# Intracellular enhancement of BMP signaling by LIM-domain protein FHL3 controls spatiotemporal emergence of the neural crest driven by WNT signaling

**DOI:** 10.1101/711192

**Authors:** Mansour Alkobtawi, Patrick Pla, Anne H. Monsoro-Burq

**Author notes:** co-last authors (Patrick Pla, Anne H. Monsoro-Burq).

## Abstract

How multiple morphogen signals are coordinated in space and time to position key embryonic tissues remains elusive. During neural crest formation, bone morphogenetic protein (BMP), fibroblast growth factor (FGF) and WNT signaling cooperate by acting either on the paraxial mesoderm or directly on the neural border ectoderm, but how each tissue interprets this complex information remains poorly understood. Here we show that Fhl3, a scaffold LIM domain protein of previously unknown developmental function, is essential for neural crest formation by linking BMP and WNT signaling thereby positioning the neural crest-inducing signaling center in the paraxial mesoderm. During gastrulation, Fhl3 promotes Smad phosphorylation and Smad-dependent *wnt8* activation specifically in the paraxial mesoderm, thus modifying the respective mesoderm or ectoderm cell response to the extracellular BMP gradient. This ensures neural border ectoderm specification by the underlying mesoderm via non-cell autonomous WNT signaling. During neurulation, neural crest inducers activate *fhl3,* promoting BMP/Smad-dependent WNT activity required for neural crest specification. Our findings highlight how Fhl3, acting cell-autonomously, ensures a fine spatial, temporal and germ layer-specific coordination of BMP and WNT signaling at several steps of neural crest development.

**Highlights:**

- FHL3 is a novel intracellular enhancer of BMP signaling during early development.
- FHL3 ensures cross-talk between BMP and WNT signaling by Smad1-dependent wnt8 activation in the paraxial mesoderm.
- FHL3 reiterated function in paraxial mesoderm and in neural border ectoderm is essential for neural crest development at the border of the neural plate.

## Introduction

A strict spatial and temporal control of BMP activity is crucial for both early mesoderm and ectoderm patterning (Bier and De Robertis, 2015). Additional morphogen gradients further refine the pattern of each germ layer. For instance the cooperation between BMP, FGF and WNT signaling pathways activates neural crest (NC) early development (Garnett et al., 2012). How these pathways crosstalk and are integrated into a coherent cell response remains poorly understood either during NC induction by the underlying mesoderm, or during its later specification at the edge of the neural plate ectoderm (NP). NC cells (NCCs) are a multipotent population generating a vast array of derivatives in vertebrates, including neurons, glia, pigment, bone and cartilage (Douarin and Kalcheim, 1999). The NC is induced from the neural border ectoderm (NB), precisely located lateral to the NP and emerging at early gastrula stage (De Crozé et al., 2011; Plouhinec et al., 2017). Initiation of the NC gene regulatory network (GRN) during gastrulation involves the response to BMP, WNTs and FGFs activated at precise levels, either in the ectoderm or in the underlying mesoderm, and diffusing towards the future NB. Later on, during neurulation and organogenesis, reiterated BMP and WNT signals control several additional steps of the NC-GRN within the ectoderm (Pla and Monsoro-Burq, 2018). While the role of each pathway has been studied separately, potential fine-tuned and tissue-specific cross-regulations between BMP and WNT signaling remain to be understood.

In the gastrula ectoderm, lack of BMP signaling allows neural induction, intermediate levels of BMP signaling are required for NB and NC initiation, while high BMP activity promotes non-neural ectoderm formation (De Robertis and Kuroda, 2004; Faure et al., 2002; LaBonne and Bronner-Fraser, 1998; Luo et al., 2001). In gastrula mesoderm, paraxial and intermediate progenitors receive moderate BMP activity due to BMP antagonists diffusing from the axial mesoderm (Harland and Gerhart, 1997). During neurulation, enhanced BMP signaling around the NP is necessary for further NC development (Litsiou et al., 2005; Steventon et al., 2009; Wu et al., 2011). Therefore, a fine-tuned and dynamic modulation of BMP signaling is particularly critical for positioning the NC, between neural and non-neural ectoderm, and promoting sequential steps of the NC-GRN. BMP ligands bind to membrane receptors triggering intracellular phosphorylation and dimerization of the receptor-regulated (R) Smad1/5/8, and their trimerization with the common (co) Smad4 (Lönn et al., 2009). R-Smad/co-Smad complexes regulate target genes transcription in the nucleus, in a BMP dose-dependent manner (Massagué et al., 2005). Numerous extracellular or cytoplasmic regulators of BMP signaling act in various cell contexts (Kuriyama et al., 2006; Schille et al., 2016; Sieber et al., 2009). During NC development however, apart from SNW1 and SMURF1, two BMP modifiers acting in the neurectoderm (Piacentino and Bronner, 2018; Wu et al., 2011), little is known about the regulators controlling the intermediate levels of BMP signaling locally early on.

Moreover, modulating BMP signaling to reach intermediate levels of activity is not sufficient to elicit robust NB induction: additional WNT and FGF signals from the mesoderm cooperate with BMPs to activate NB/NC specifier expression in the ectoderm. At the time of NB specification in frog gastrulas, the dorsal-lateral marginal zone (DLMZ), which will form both paraxial and intermediate mesoderm (thereafter collectively called “paraxial mesoderm”), expresses several FGF and WNT ligands diffusing towards the overlying ectoderm. Inhibiting any of these signals prevents NB-NC activation (Chang and Hemmati Brivanlou, 1998; Elkouby et al., 2010; Garnett et al., 2012; Garriock et al., 2007; LaBonne and Bronner-Fraser, 1998; Monsoro-Burq et al., 2003; Saint-Jeannet et al., 1997).

We found enriched expression of *fhl3,* encoding an intracellular scaffolding protein, in the developing frog NC and underlying mesoderm. Fhl3 is a four-and-a-half LIM domain protein implicated in protein-protein interactions in adult cells and cancer, with no previously described developmental function (Kadrmas and Beckerle, 2004). FHL proteins broadly regulate cell proliferation, differentiation and apoptosis (Bach, 2000). In cancer cells responding to TGFβ, Fhl3 interacts with R-Smad2/3 and Smad4 and modulates TGFβ signaling in a context-dependent manner (Ding et al., 2009; Han et al., 2018). During muscle fiber differentiation *in vitro*, Fhl3 binds to MyoD and to Sox15, resulting in *Foxk1* coactivation, increased myoblast proliferation and blocked myotube fusion (Cottle et al., 2007; Meeson et al., 2007). These studies thus suggest that Fhl3 could modulate both cell signaling or gene transcription.

Here, we find that Fhl3 is a critical enhancer of BMP and WNT signaling during NC early development *in vivo*, linking the BMP intracellular signaling cascade to *wnt8* activation both in the mesoderm and in the neural border ectoderm. Fhl3 cell-autonomous activity allows modulating independently mesoderm or ectoderm cell response to the same extracellular BMP gradient and is essential for the coordinated patterning of dorsal tissues during NC development.

## Results

### Fhl3 is expressed in the dorsal mesoderm and activated in neural crest cells by Pax3

To get insights into *fhl3* function, we analyzed its developmental expression in frogs, compared to patterns of regional markers in the dorsal mesoderm and ectoderm (Figures 1A, S1). *Fhl3* expression appears in the involuting mesoderm of early gastrulas (dorsal marginal zone (DMZ) and DLMZ) and remains in the axial (notochord) and paraxial mesoderm until tailbud stages. *Fhl3* is also expressed in the late gastrula NB ectoderm (Figure S1). In mid-neurula ectoderm, when compared to the NB marker *pax3* and the premigratory NC marker *snai2, fhl3* is first activated in a subset of cephalic NCCs at hindbrain level. Upon neural tube closure, *fhl3* is expressed in all premigratory NCCs. In tailbud-stage embryos *fhl3* is detected in the NCCs migrating towards the branchial arches with a stronger expression in the hyoid stream, compared to *twist1* which labels all migrating NCCs. In tadpoles, *fhl3* is observed in branchial arches, notochord and somites. In summary, *fhl3* was detected in two key tissues involved in NC formation: in the dorsal mesoderm, at the time of NB induction by the underlying paraxial mesoderm, i.e. upstream of NC specification, and later on, in the premigratory, migrating and differentiating cranial NCCs themselves.

**Figure 1.**
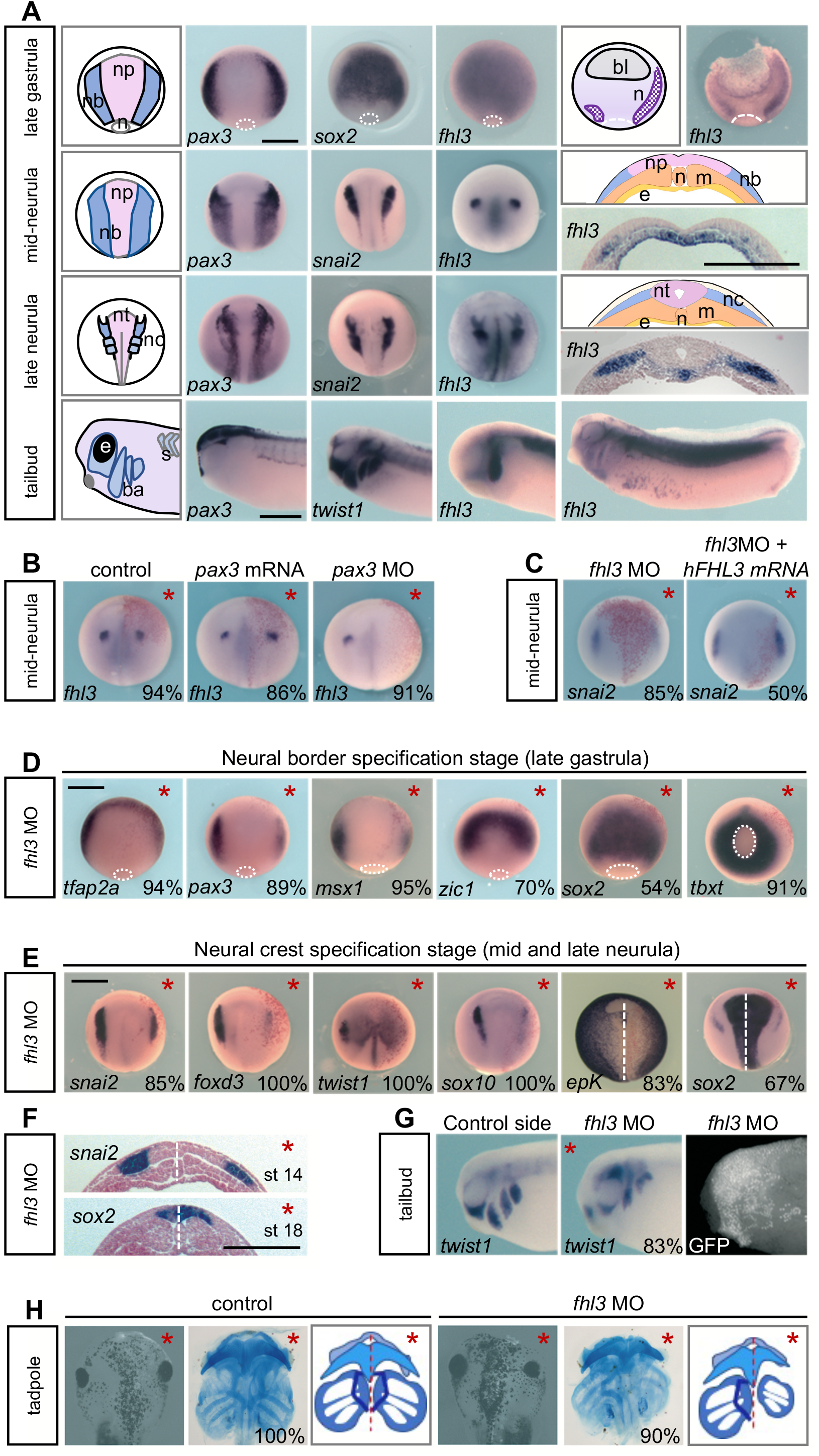
Fhl3 is required for NB and NC induction. **(A)** Comparison of *fhl3* expression to neural, neural border and NC markers during frog development. np, neural plate; nb, neural border; nt, neural tube; n, notochord; bl, blastocoel; n, neural crest; m, paraxial mesoderm; e, endoderm; ba branchial arches; s, somite. **(B)** Pax3 gain (mRNA) or loss of function (MO) regulates *Fhl3* expression *in vivo*. **(C)** Human *FHL3* rescues the *fhl3* morphant phenotype *in vivo*. (**D-F)** *Fhl3* depletion prevents neural border (D, F) and neural crest (E, F) specifiers gene activation but not neural and mesoderm markers expression. **(G-H)** *Fhl3* depletion blocks NC migration and cranio-facial cartilage formation. Scale bar: 500 μm.

Interestingly, *fhl3* expression partly overlapped with *pax3* which is essential for NC induction in the ectoderm, consistent with the identification of *fhl3* as a putative target of Pax3 (Bae et al., 2014, and our unpublished data). To test the role of Pax3 in *fhl3* induction in NC, we either increased or depleted Pax3 unilaterally, using mRNA injections or a validated morpholino oligonucleotide (MO, Monsoro-Burq et al., 2005). *Fhl3* expression was modestly expanded upon Pax3 gain of function and eliminated after Pax3 depletion *in vivo* (Figure 1B, S3).

#### Fhl3 is essential for neural border specification in gastrulas and neural crest induction in neurulas

To assess its developmental function *in vivo*, we depleted Fhl3 on one side of frog embryos using either a translation-blocking MO, or a splice-blocking MO or CRISPR/cas9-mediated *fhl3* mutation (Figures 1C-H, S2–S3). MO specificity and lack of toxicity were assessed by a phenotype rescue with human FHL3, also indicating the evolutive conservation of the protein’s function (Figure 1C). After Fhl3 depletion, NB markers *tfap2a*, *pax3* and *msx1* were severely downregulated on the injected side of late gastrulas, while others such as *zic1* were unchanged or even upregulated (Figure 1D). Expression of the NP marker *sox2* was expanded. This demonstrated that *fhl3* morphant NB tissue was present but mispatterned. Expression of the pan-mesoderm marker *tbxt* (*xbra*) or of the paraxial mesoderm marker *myod* were unaffected indicating globally normal mesoderm formation. These results indicate that Fhl3 does not affect the global formation of dorsal mesoderm and ectoderm during gastrulation but is essential for NB early patterning.

During neurulation, Fhl3 knockdown caused the downregulation of all premigratory NC markers tested: *snai2, foxd3, twist1* and *sox10* (Figure 1E). Instead, *sox2* expansion indicated that the NP was enlarged, at the expense of NC and non-neural ectoderm (marked by *epidermal keratin epK).* Transverse sections confirmed the lateral expansion of the neural tissue (*sox2*) and the lateral shift of the reduced *snai2* territory (Figure 1F). At tailbud stage, Fhl3 morphant cranial NCC migration into the pharyngeal arches was markedly reduced (labeled by *twist1* expression, Figure 1G), resulting in severe alterations of craniofacial skeleton differentiation (Figure 1H). In particular, the *trabeculae cranii* derived from the mandibular and hyoid arches were missing on the injected side and branchial arches were reduced. Together, these results indicate that during NC formation, Fhl3 is necessary for NB and NC induction and that the early NC developmental defects caused by Fhl3 depletion were not compensated at later stages of NC differentiation.

#### Fhl3 controls BMP signaling in the paraxial mesoderm

The altered gene patterns observed above after Fhl3 depletion could result from abnormal signaling events in the early NB/NC-GRN. Lower BMP signaling could expand the neural plate and shift/reduce the NB laterally. During NB induction, *zic1* expression is triggered by low BMP signaling, while *tfap2a*, *pax3,* and *msx1* are activated by high WNT signaling (De Crozé et al., 2011; Mizuseki et al., 1998). Thus, low BMP and low WNT signals could result in the observed gene pattern deregulations. Since Fhl3 was previously shown to mediate TGFβ signaling in cancer cells, we asked whether the early loss of Fhl3 affects BMP activity in the prospective paraxial mesoderm. A BMP signaling reporter driving luciferase expression was injected into embryos together with *fhl3* MO. The DLMZ was dissected out of early gastrulas (Figure 2A). Luciferase activity was significantly reduced in *fhl3* morphant DLMZ compared to controls (Figure 2B). Furthermore, phosphorylation of R-Smad1-5-8 (pSmad1/5/8) was strongly decreased in *fhl3* morphant DLMZ (Figure 2C). Both results demonstrate that Fhl3 depletion strongly lowers BMP signaling in the paraxial mesoderm.

**Figure 2.**
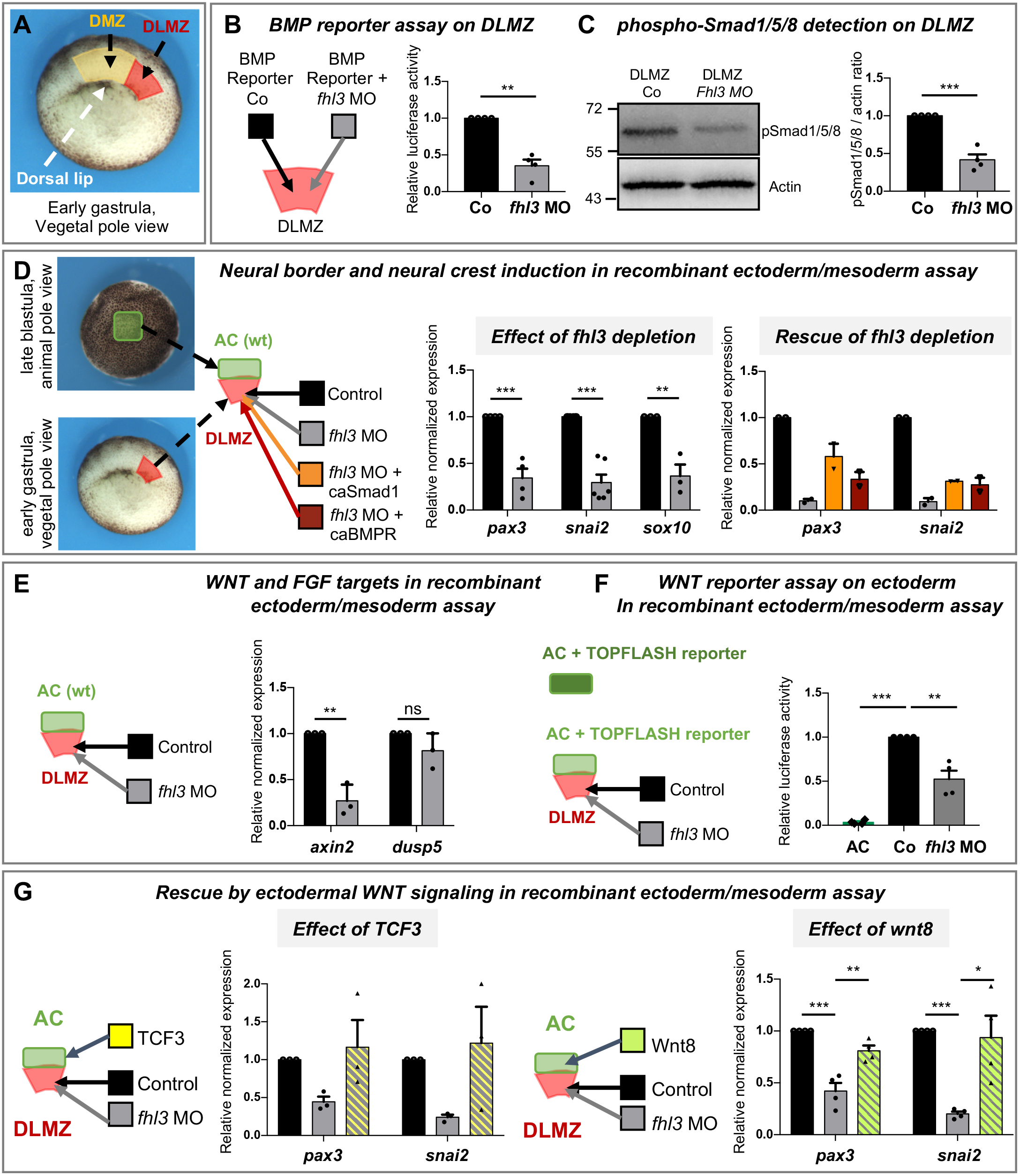
Fhl3 is necessary for BMP signaling and WNT-dependent NC inducing activity in the mesoderm. **(A-C)** Fhl3 is essential for BMP activity levels and Smad1/5/8 phosphorylation in the paraxial mesoderm of late-gastrulas. **(D)** In mesoderm (DLMZ)/ectoderm (AC) NC-inducing assay, depleting Fhl3 selectively in the mesoderm affects NB/NC induction in the ectoderm. This phenotype is partially rescued by the co-injection of *ca-smad1* or *caBMPR* in the mesoderm. **(E)** In the same assay, both *axin2* expression (WNT target) and WNT reporter activity are decreased when ectoderm is juxtaposed to *fhl3* morphant mesoderm. In contrast, *dusp5* expression (FGF target) remains unchanged. **(F)** WNT reporter TOPFLASH-injected ectoderm showed decreased WNT response when juxtaposed to a *fhl3* morphant DLMZ, compared to a control DLMZ. **(G)** Injection of either *tcf3* or *wnt8* mRNAs in the ectoderm rescues NC induction in ectoderm juxtaposed to *fhl3* morphant mesoderm.

To investigate the impact of Fhl3 depletion on the NB-NC-inducing ability of the paraxial mesoderm, we used a classical assay in *X. laevis* embryos (Bonstein et al., 1998): the recombination of a DLMZ, dissected from an early gastrula, with a fragment of pluripotent blastocoel roof ectoderm dissected from a late blastula (animal cap, AC) (Figure 2D). When compared to control DLMZ, *fhl3* morphant DLMZ was unable to activate NB and NC markers *pax3*, *snai2* and *sox10*. Moreover, to test if the altered BMP signaling observed in the morphant mesoderm was causing its lack of inducing activity, we attempted a phenotype rescue by co-injecting fhl3 MO with a constitutively active form of Smad1 (Smad1 3SD) or of BmpR1 (caBMPR). In this case, the co-injected DLMZ recovered ability to activate *pax3* and *snai2* expression albeit with lower efficiency compared to uninjected control. These results demonstrate that Fhl3 mesodermal function is essential for NC induction in the ectoderm by the paraxial mesoderm, at least partly by promoting BMP signaling.

#### Fhl3 controls vertical Wnt8 signaling acting between paraxial mesoderm and ectoderm during NC induction

Wnts and FGFs from the paraxial mesoderm are crucial for inducing the NC. Therefore, we quantified the expression of *dusp5,* a FGF-responsive gene, and *axin2*, a readout for canonical WNT signaling, in mesoderm-ectoderm recombinants (Figure 2E). *Axin2* expression, but not *dusp5* levels, was reduced in the *fhl3* morphant recombinants suggesting altered WNT activity and normal FGF function. To confirm that Fhl3 depletion in the mesoderm resulted in altered canonical WNT signaling specifically in the ectoderm, we injected a WNT/β-catenin signaling reporter vector (TOPFLASH) into the ectoderm only and juxtaposed it to a DLMZ dissected either from uninjected or from *fhl3* morphant embryo. Compared to the control situation, the ectoderm combined to morphant mesoderm showed a significantly decreased response to WNT signaling (Figure 2F). These results thus show that the levels of WNT activity in the ectoderm depend on Fhl3 function in the DLMZ.

To test if the decreased WNT signaling observed in the ectoderm following Fhl3 mesodermal depletion was sufficient to explain the lack of NC induction in the ectoderm, we implemented a phenotype rescue using either an inducible form of activated TCF3 or *wnt8* gain-of-function, injected in the ectoderm only (Figure 2G). When uninduced ectoderm was juxtaposed to Fhl3-depleted DLMZ, *pax3* and *snai2* expression levels were low, as observed with uninjected ectoderm previously. However, upon induction of TCF3 or increased Wnt8 expression in the ectoderm, *pax3* and *snai2* levels were fully recovered. Together, these results demonstrate that Fhl3 early function in the paraxial mesoderm is critical for its ability to activate the canonical WNT signaling at levels required for NC induction in the ectoderm.

#### During neurulation, Fhl3 is necessary for NC induction by neural border specifiers and controls both BMP and WNT signaling levels in the ectoderm

During the subsequent steps of the NB/NC-GRN, *fhl3* is activated by Pax3 in the NC cells (Figure 1). In order to separate the roles of *fhl3* in NC formation within the ectoderm from its functions in the mesoderm, we have used direct NC induction in pluripotent blastocoel roof ectoderm (induced NC, iNC) by co-injecting calibrated levels of Pax3 and Zic1, a combination of two transcription factors necessary and sufficient for NC induction and development from the early ectoderm (Milet et al., 2013, Figure 3A). Compared to control iNC, co-injecting *fhl3* MO prevents the activation of NC markers *snai2* and *sox10* suggesting that Fhl3 is also required for premigratory NC specification downstream of NB regulators (Figure 3B). Moreover, Pax3 displays a positive autoregulatory loop in the ectoderm important for *pax3* maintenance in the NB (Plouhinec et al., 2014). Fhl3 morphant ectoderm seems defective for this autoregulation, since *pax3* expression is decreased as well.

**Figure 3.**
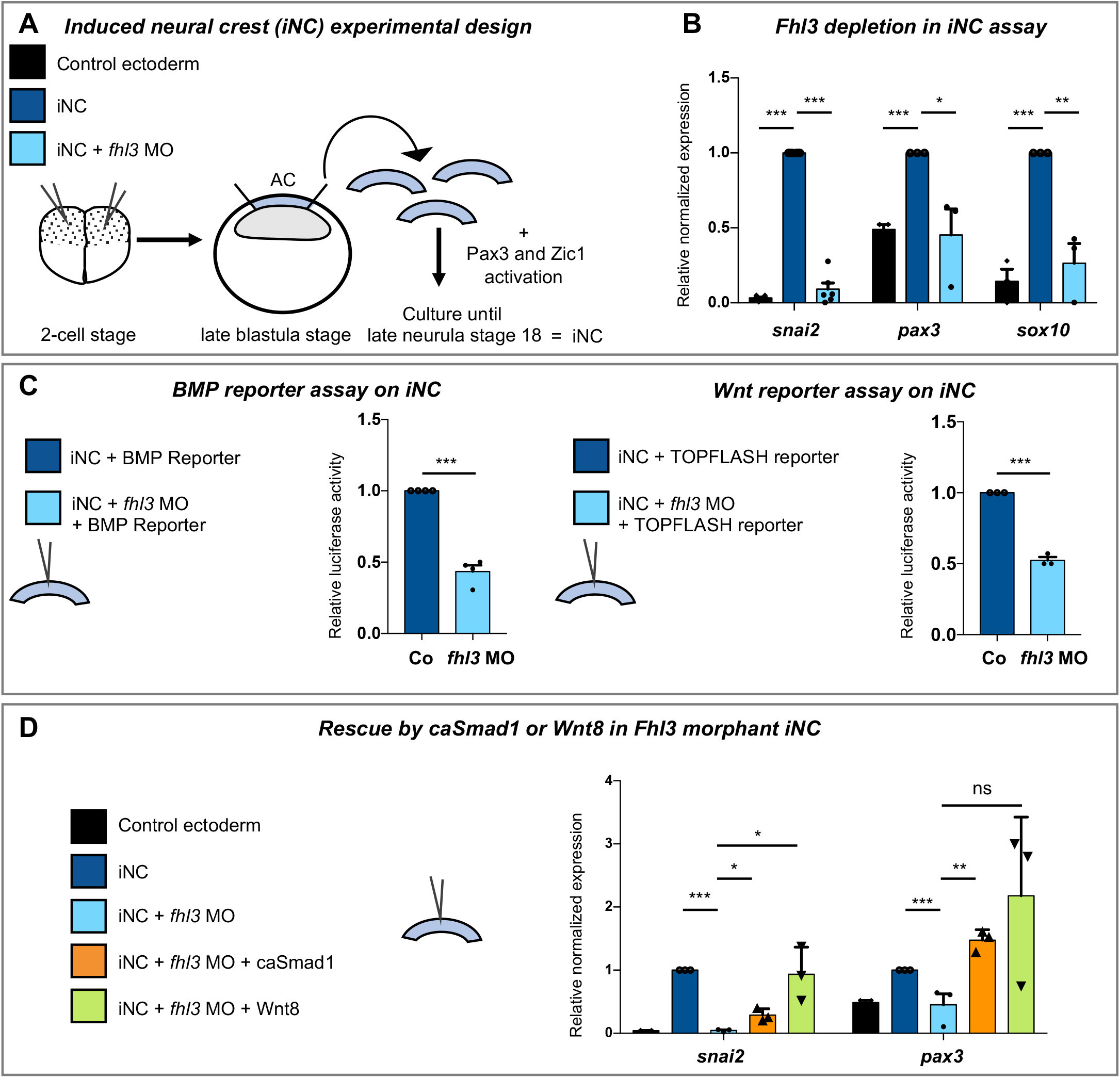
Fhl3 is necessary for NC specification in the ectoderm, downstream of neural border specifiers Pax3 and Zic1. **(A-B)** In a direct NC induction assay, key NC specifiers are poorly induced in *fhl3* morphant iNC compared to control iNC. **(C)** BMP and WNT activities are reduced in *fhl3* morphant iNC. **(D)** Injection of *ca-smad1* or *wnt8* rescue NC induction in *fhl3* morphant iNC.

Next, we tested whether Fhl3 depletion in iNC affected BMP and WNT signaling as observed in the mesoderm. When either the BMP or the WNT signaling reporters were co-injected with Pax3 mRNA, Zic1 mRNA and *fhl3* MO, the luciferase activity was severely reduced compared to signal in control iNC (Figure 3C). Finally, co-injection of either caSmad1 or *wnt8,* in *fhl3* morphant iNC were sufficient to rescue *snai2* and *pax3* expression (Figure 3D). Altogether, these data indicate that, during neurula-stage steps of the NC-GRN, Fhl3 promotes both BMP and WNT signaling in the ectoderm and that this regulation is necessary to achieve NC induction.

#### Smad1 and Fhl3 directly bind *wnt8* promoter

The parallels between BMP activity and WNT signaling observed in the series of results described above both in the mesoderm and in the ectoderm during NC development, led us to hypothesize that Fhl3 could mediate a cross-talk between the two signaling pathways. In Fhl3 morphant mesoderm, we observed diminished expression of *wnt8* and WNT-responsive gene *myf5* on the injected side (Figures 4A, S2C). In addition, the decreased pSmad1/5/8 levels suggested a post-translational regulation downstream of BMPR (Figure 2C), while the decreased *wnt8* expression suggested a transcriptional regulation. In line with this hypothesis, we found Smad binding elements in the proximal *wnt8* promoter region (BS1-2, Figure 4A). To examine Smad1 occupancy at these sites, we injected tracing amounts of FLAG-tagged Smad1 in whole embryos and used an anti-FLAG antibody at late gastrula stage for *in vivo* chromatin immunoprecipitation followed by quantitative PCR (ChIP-qPCR): strong Smad1 occupancy was detected at both sites (binding to BMP-responsive gene *ventx1* serves as a positive control, Figure S2F). When the embryos were injected with FLAG-tagged Fhl3, clear Fhl3 binding was observed on the same BS1-2 on *wnt8* promoter, showing that Smad1 and Fhl3 are likely engaged together in the transcriptional complex regulating *wnt8* expression (Figure 4B). When BMP signaling was decreased by injecting the extracellular BMP antagonist Noggin in whole embryos, Fhl3 signal was reduced, showing that Fhl3 specifically binds these sequences in response to active BMP signaling (Figure 4C). Finally, after Fhl3 depletion either in the mesoderm or in the NC-induced ectoderm, gain-of-function of caSmad1 or caBMPR restores *wnt8* and *axin2* expression, indicating that the function of Fhl3 in the NC-GRN is to enhance Smad1 activity which in turn up-regulates *wnt8* expression (Figure 4D). In sum, Fhl3 is a novel and key coordinator of BMP/Smad1 and WNT/β-catenin signaling that tunes the levels of activity of these pathways in dorsal tissues to trigger NC induction during the two first steps of the NC-GRN (Figure 4E).

**Figure 4.**
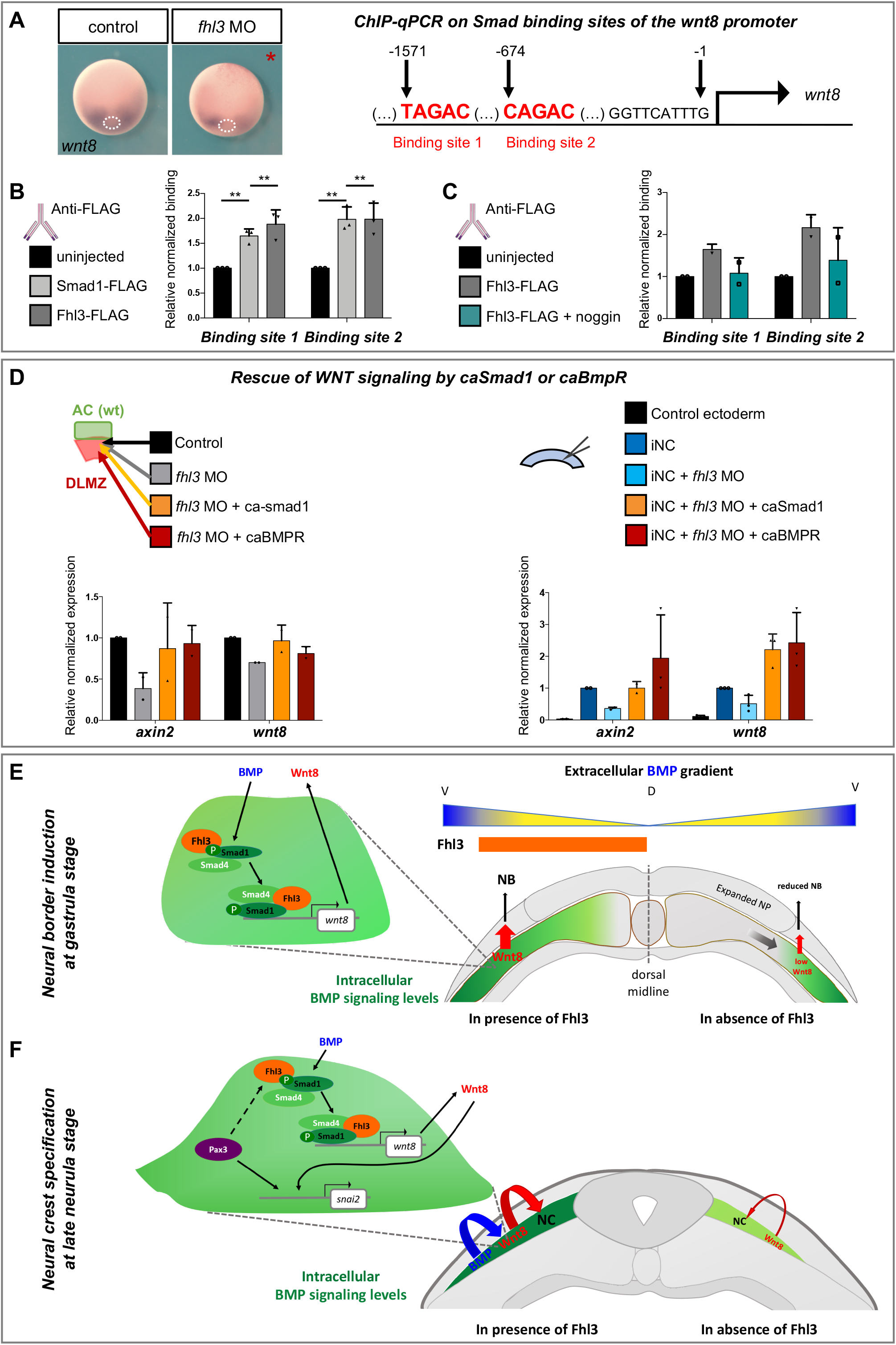
Fhl3 enhances Smad1-dependent *wnt8* transcription. **(A)** *Wnt8* expression is decreased in *fhl3* morphant mesoderm *in vivo*. **(B-C)** At late gastrula stage, ChIP-qPCR analysis reveals binding of both Smad1 and Fhl3 on two Smad binding sites found in *wnt8* promoter (BS, shown in red). When extracellular BMP signaling is antagonized by Noggin, Fhl3 binding on BS is strongly reduced. **(D)** *Wnt8* and *axin2* expression are rescued by *ca-smad1* or *caBMPR* either in *fhl3* morphant recombinants or in morphant iNC. **(E, F)** Model of the sequential roles of Fhl3 at two critical steps of NB and NC induction. Combined activity of the ventral-dorsal BMP gradient (blue/high to yellow/low) and of dorsally expressed Fhl3 (orange) positions the NC-inducing signaling center in the paraxial mesoderm during gastrulation (E). (F) During neurulation, Fhl3 enhances BMP and WNT signaling in the premigratory NC enhancing NC specification.

## Discussion

Early patterning and later organogenesis are the result of a remarkably orchestrated suite of signaling and inductive interactions. Here, we used time-controlled and tissue-specific gene manipulations combined to embryonic tissue recombinations to disentangle the sequence of events involving Fhl3 in early NB/NC induction. We find that early gastrula dorsal mesoderm simultaneously expresses BMP modulators with opposing activities: the known secreted BMP antagonists diffusing in the extracellular space and blocking BMP signaling both in ectoderm and mesoderm, and a novel BMP signaling agonist, FHL3, acting cell-autonomously in the mesoderm. Consequently, paraxial and intermediate mesoderm cells, although receiving low BMP signals, maintain an intracellular phospho-Smad1/5/8 activity sufficient to directly activate Wnt8 signaling (Figure 4E). In the absence of Fhl3, the initial events of mesoderm patterning are intact (tissue morphology, *tbxt* expression, Figure 1) but the downstream mesoderm signaling activity during gastrulation is impaired (BMP, WNT, Figure 2). As a result, WNT-responsive NB specifiers are not activated and NB ectoderm induction fails and the neural plate is expanded, ultimately preventing NC development (Figure 1, 4). This is the first demonstration of a molecular link coordinating BMP and WNT signaling at the root of the NB/NC-GRN during gastrulation. Furthermore, during neurulation, WNT signals reiteratively cooperate with NB transcription factors such as Pax3, Tfap2a and Zic1, to activate the premigratory NC program (Monsoro-Burq et al., 2005; Sato et al., 2005; Simões-Costa et al., 2015). In particular, *snai2* is directly activated by both Pax3 and WNT signals (Plouhinec et al., 2014; Vallin et al., 2001). Here we show that Pax3 also activates *fhl3* in the prospective NC, and that Fhl3 promotes WNT signaling in a Smad-dependent manner in premigratory NC cells (Figures 1, 3, 4D). This creates a positive amplification loop, encoded within the NC-GRN and coordinating both secreted and transcriptional NC inducers to achieve a robust NC specification (Figure 4F). To conclude, Fhl3 is a novel regulator of BMP signaling, modulating the spatial and temporal response to BMPs in a germ layer-specific manner, and coordinating BMP and WNT signaling at several stages of NC development. In the global view of early development, our findings highlight how regional patterns are achieved in vertebrate embryos following the initial dorsal-ventral patterning: the cooperation between dorsal (Fhl3) and ventral (BMP) cues positions a secondary signaling center in the paraxial mesoderm, sending WNT signals towards an ectoderm area receiving low extracellular BMP signaling, ultimately inducing the neural crest.

## Acknowledgments.

We are gratefult to Drs K.J. Liu and E. Theveneau for their insightful comments on the manuscript, to all members of the Monsoro-Burq team especially M. Sittewelle and P. Scerbo for proofreading the manuscript. We also thank S. Dodier for her technical help with histological sections, E. Belloir for animal husbandry and the Korean Xenopus Resource Center for Research (XRCR-Hallym University, Korea) for providing constructs.

This work was funded by Université Paris Sud, Centre National de la Recherche Scientifique (CNRS), Agence Nationale pour la Recherche (ANR-15-CE13-0012-01-CRESTNETMETABO), Fondation pour la Recherche Médicale (DEQ20150331733) and Institut Universitaire de France to AHMB. M. Alkobtawi is a Ph.D. fellow funded by Fondation pour la Recherche Médicale (DEQ20150331733) and Institut Curie.

---

**Supplementary Figure S1.**
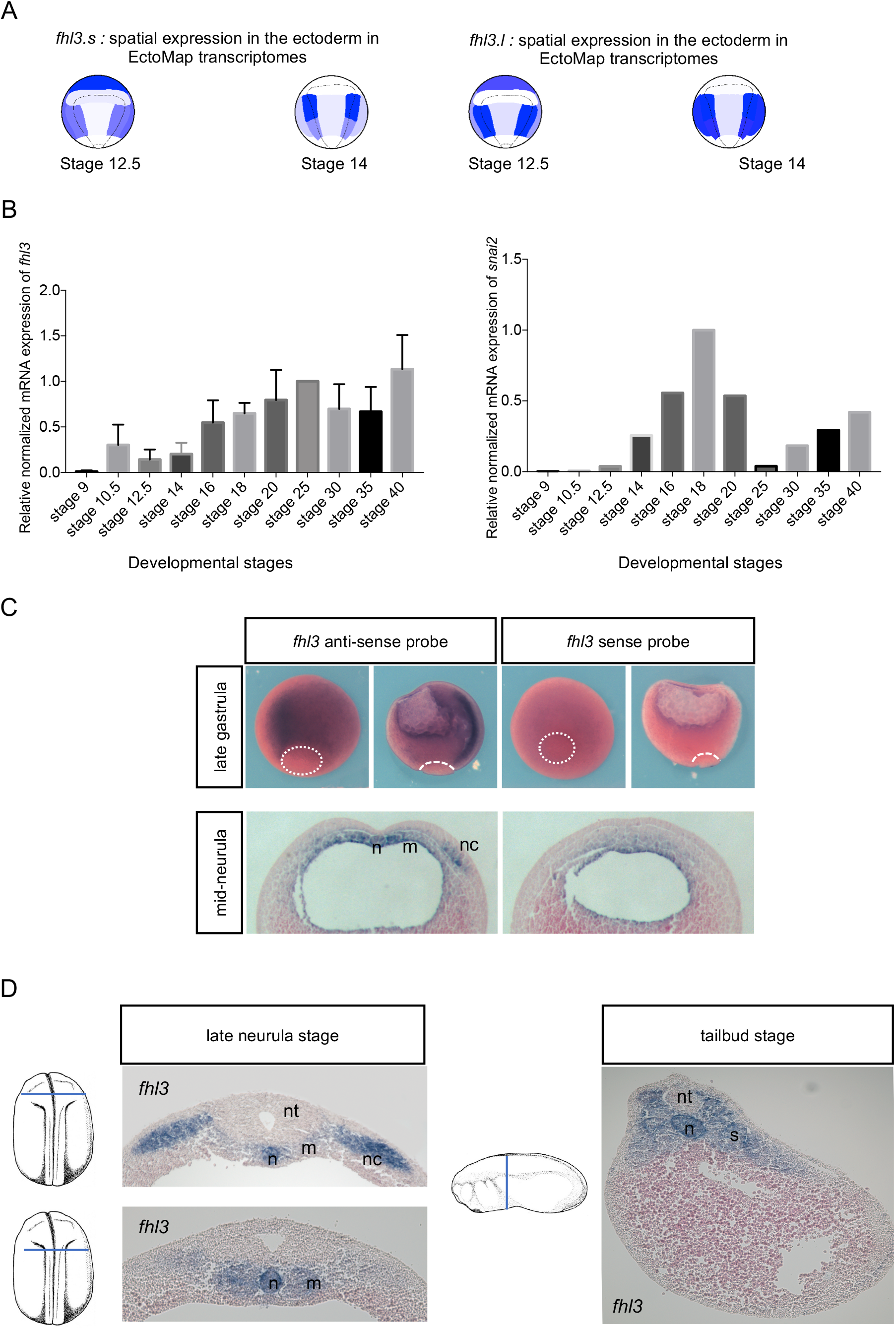
*fhl3* is detected in the early involuting mesoderm, the neural border ectoderm, neural crest cells, notochord and somites. **(A)** Ectoderm-specific regional transcriptome data sets (EctoMap) predict *fhl3.s* and *fhl3.l* expression in the NB ectoderm and in the non-neural ectoderm at lower levels, at late gastrula and mid-neurula stages (Plouhinec et al., 2017). **(B)** qRT-PCR analysis on whole embryos finds the earliest expression of *fhl3* at early gastrulation (stage 10.5), prior to *snai2* onset (late gastrula stage 12.5). Strong *fhl3* expression is observed from neurula (stage 16) to tadpole stages (stage 40). **(C)** WISH and sagittal sections after staining with anti-sense or sense probes for *fhl3* show the specificity of the staining with the antisense probe in the involuting mesoderm at early and late gastrula stages (stage 10.5 and stage 12.5). Sections further show the staining specificity in the NC, the axial and the paraxial mesoderm. **(D)** Transverse sections at late neurula stage show *fhl3* expression in the premigratory NC, the notochord and the somites. Transverse sections at tailbud stage show *fhl3* expression in the notochord and the somites. nt, neural tube; n, notochord; nc, neural crest; m, paraxial mesoderm; s, somites.

**Supplementary Figure S2.**
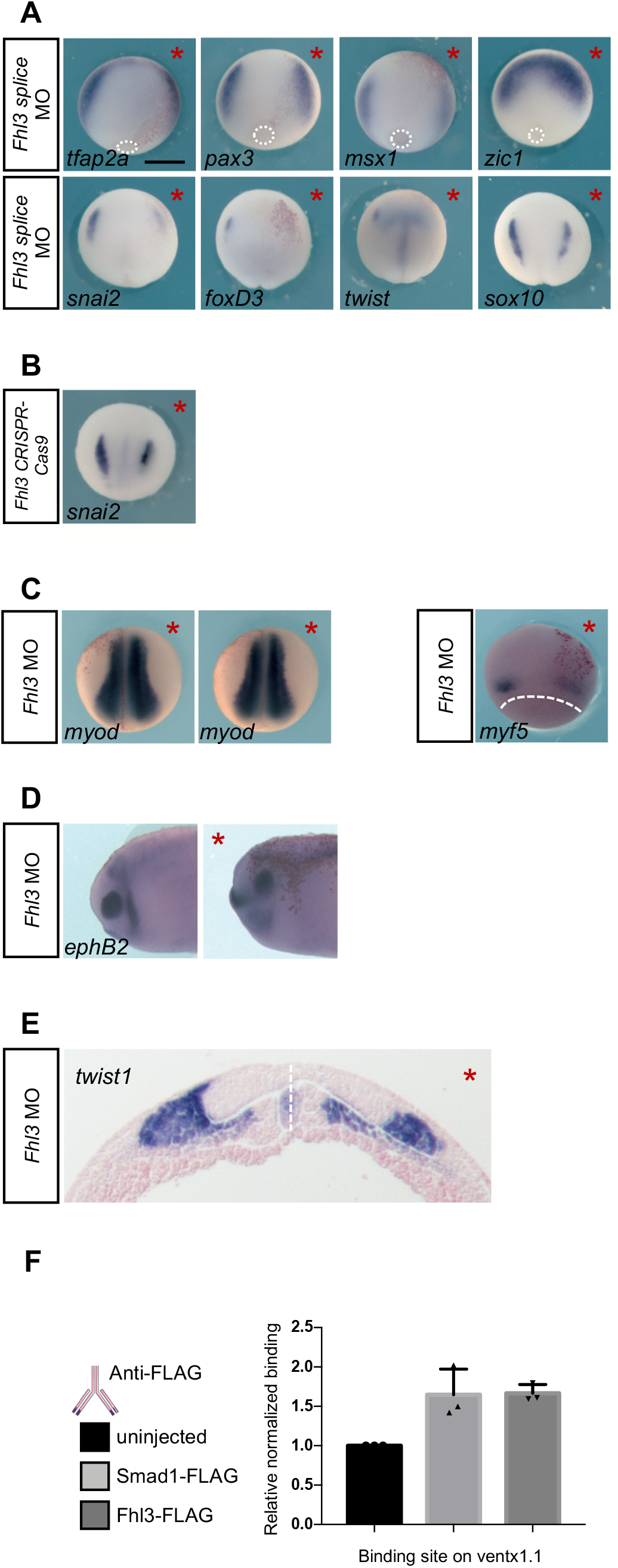
*fhl3* mutation by Crispr-cas9 and a *fhl3* depletion using a splice-blocking MO reproduced the *fhl3* ATG-MO phenotypes on neural crest and mesoderm. **(A)** A *fhl3* splice-blocking MO (SP-MO) reproduces the *fhl3* ATG-MO phenotype. Expression of the NB markers *tfap2a*, *pax3* and *msx1* is decreased in *fhl3* SP-MO. NB marker *zic1* is unaffected or slightly increased. Similarly, NC markers *snai2, foxd3*, *twist1* and *sox10* expression is decreased in *fhl3* SP-MO. **(B)** Injection of *fhl3* gRNA and Cas9 protein into one blastomere of two-cell stage embryos decreases *snai2* expression at mid-neurula stage. The injected side is indicated by *. **(C)** Fhl3 depletion does not affect the expression of the paraxial mesoderm marker *myod* (late neurula stage) but decreases the expression of the WNT-responsive gene *myf5* at (gastrula stage). **(D)** At tailbud stages, the expression of *ephB2,* a second branchial arch maker, is lost in *fhl3* morphant embryos. The injected side is indicated by *. **(E)** In transverse sections, *twist1* expression is shifted laterally and reduced in the injected ectoderm (*) compared to the control side, while *twist1* expression boundaries in the mesoderm remain unaffected by Fhl3 depletion on the injected side. Dashed white line indicates the midline.

**Supplementary Figure S3.**
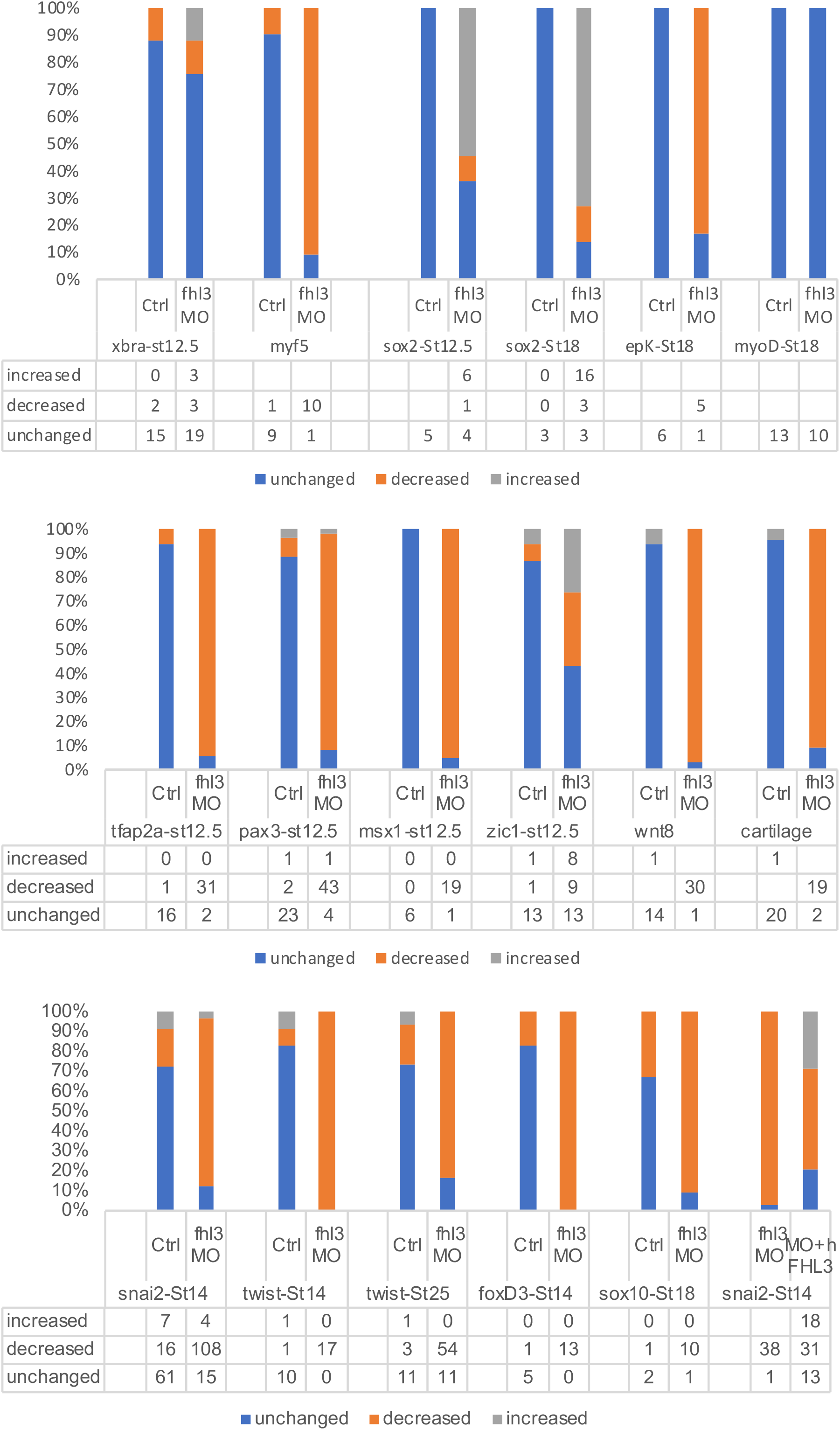

**Supplementary Figure S4.**
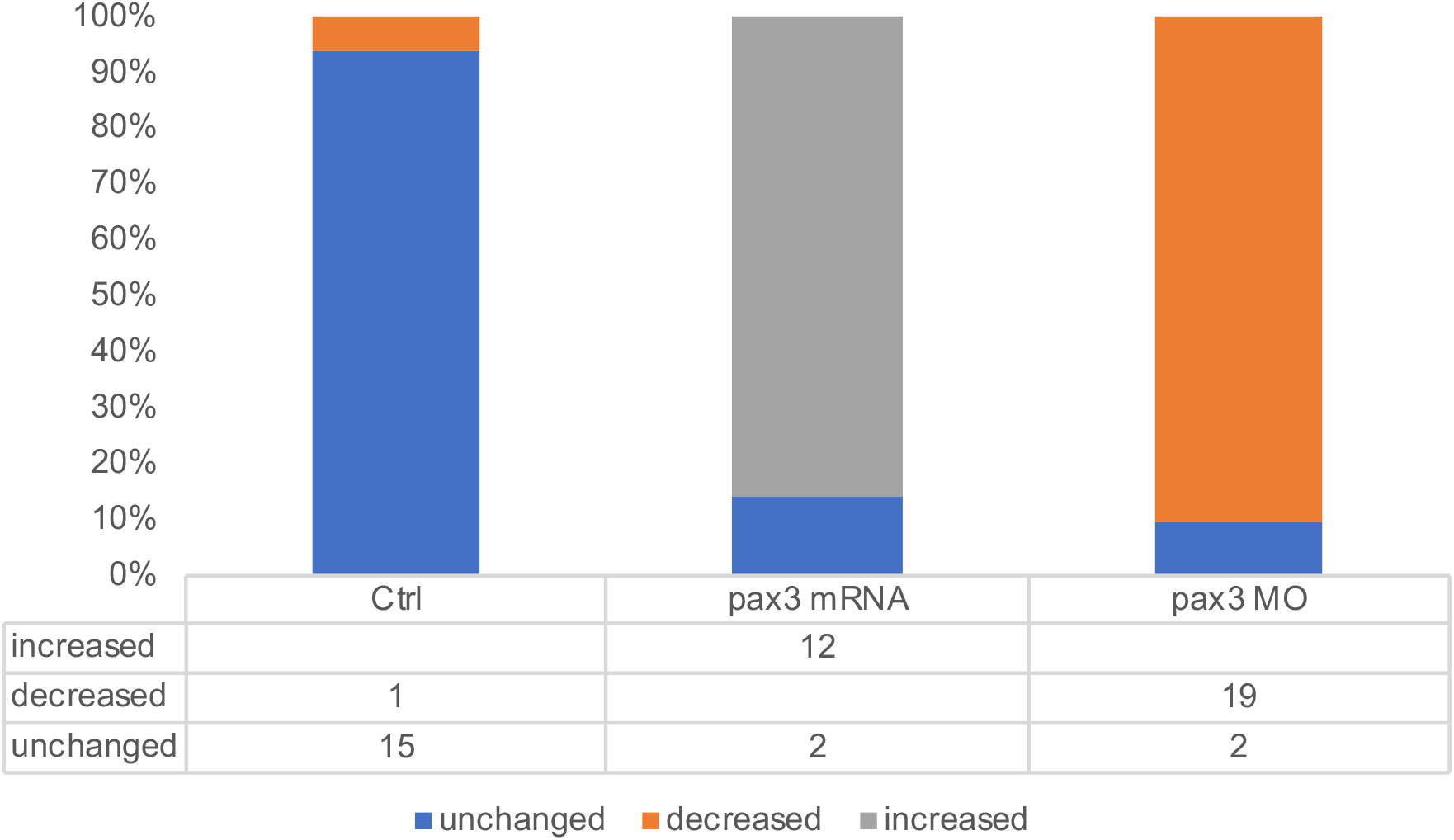

## Materials and Methods

### Sub-cloning of *Xenopus laevis fhl3* cDNA

An *fhl3* expression vector was obtained by amplifying the full-length *fhl3* cDNA by PCR from late neurula stage whole embryo cDNA, and inserting it into pGEM-T Easy (Promega) by TA cloning. The insert was then subcloned into pCS107 (*Not*I site). *Fhl3* PCR primers are listed in Table S1.

### Xenopus embryos manipulations

Ovulation, *in vitro* fertilization and embryo culture were performed using standard methods described in Sive et al., (2000). Embryos were staged according to P.D. Nieuwkoop and J. Faber (1995). All procedures were performed according to the European animal use and care law (animal care and housing approval license #C91-471-108, Direction Départementale de Protection de la Population, Courcouronnes, France).

### *In vivo* gene depletion, gain of function and rescue experiments

For knockdown experiments, two antisense morpholino oligonucleotide (MO) were synthesized to block *fhl3* translation or splicing (Gene Tools, Table S1). 15 ng of control MO or *fhl3* MO (either ATG MO or splice MO) were injected into one or two blastomeres of two-cell stage embryos, depending on the assay. For Pax3 depletion experiments, 20 ng of previously validated MO (Monsoro-Burq et al., 2005) was injected into one blastomere at two-cell stage embryo.

Capped mRNAs were synthesized using mMessage mMachine kits (Invitrogen). For Pax3 gain of function experiments, *pax3* mRNA (75 pg) was injected into one blastomere at two-cell stage embryo. For rescue experiments, h*FHL3* mRNA (300 pg) was co-injected with the *fhl3* MO into one blastomere of two-cell stage embryos and caSmad1 (3SD) mRNA (200 pg) or caBMPR1 mRNA (200 pg) or TCF3 mRNA (80 pg) or Wnt8 mRNA (100 pg) were co-injected with the *fhl3* MO into both blastomeres of two-cell stage embryos.

In each case, blastomeres were co-injected with β-gal mRNA (250pg) or GFP mRNA (200pg) to identify the injected side (marked by a red asterisk).

For CRIPSR/cas9 experiments, one sgRNA primer was synthesized at the Cold Spring Harbor Laboratory during the Cell and Developmental Biology of Xenopus Course. We synthesized *fhl3* gRNA using MEGAshortscript T7 transcription kit (Table S1). Cas9 protein (2.5 ng) and *fhl3* sgRNA (300 pg) were co-injected into one blastomere of two-cell stage embryos.

### Neural crest induction in mesoderm/ectoderm recombination assay

To recombine mesoderm at NC-inducing stage to responsive “naive” ectoderm, a heterochronic recombination was used (Bonstein et al., 1998). Animal caps were dissected out from blastula stage embryos (stage 9). Simultaneously, dorsal-lateral marginal zone (DLMZ) explants were dissected out from gastrula stage embryos (stage 10.5), defined when a faint pigment line is formed at the dorsal lip by the involuting cells. Recombinants were formed by juxtaposing the two tissues to one another, and cultured in ¾ NAM medium (Slack and Forman, 1980) until sibling embryos reached the late neurula stage 18.

### Direct neural crest induction (iNC)

Animal cap ectoderm is a source of pluripotent cells that respond to Pax3/Zic1 activation by a complete NC development (Milet et al., 2013). For such iNC (induced Neural Crest) assays, *pax3-GR* mRNA (35 pg) and *zic1-GR* mRNA (72 pg) synthesized with mMessage mMachine kits (Invitrogen), were co-injected into both blastomeres of two-cell stage embryos, with or without 15 ng *fhl3* MO. For rescue experiments, *caSmad1* (200 pg) or *wnt8* mRNA (100 pg) were co-injected with *pax3-GR* mRNA*, zic1-GR* mRNA and *fhl3* MO. Animal cap explants were dissected out at stage 9 and dexamethasone (or its solvent ethanol as a control) was added at stage 10.5. Explants were lysed at stage 18.

### Whole mount *in situ* hybridization (WISH) and sectioning

Embryos were stained using a fast whole-mount *in situ* hybridization protocol (Monsoro-Burq, 2007). *Pax3*, *tfap2a*, *msx1* and *zic1* are expressed in the neural border and *snai2*, *foxd3*, *twist1* and *sox10* are expressed in the neural crest. *Sox2* is a general neural plate and neural tube marker and *epK* is a non-neural ectoderm marker. *Tbxt* is a general mesoderm marker*. Myod and myf5* are expressed in the paraxial-fated mesoderm.

Late gastrula (stage 12.5), mid-neurula (stage 15) and late neurula (stage 18) embryos are shown in dorsal views with anterior to the top, except for the late gastrula stage embryo stained with *tbxt* shown in vegetal view, dorsal to the top (Figure 1D). The blastopore of late gastrulas is marked by a white dashed circlet. The dotted lines in Figure 1E indicate the midline. Tailbud stage embryos are seen in side views, anterior to the left. A sagittal view of a dissected late gastrula embryo is shown in the top right panel in Figure 1A. Mid- and late neurula embryos were embedded in paraffin and 12-µm-thick microtome transverse sections were cut (Figure 1 A, F).

### Cartilage staining

Stage 45 tadpoles were fixed, dehydrated and stained in Alcian Blue for 24 hours. After several washes in ethanol until no blue stain is released, embryos were rehydrated, cleared with 4% KOH and transferred into graded glycerol solution in 2% KOH. Finally, ventral head cartilage was dissected. Whole tadpoles are shown from the dorsal side, anterior to the top and the matching dissected cartilages are viewed from the ventral side, anterior to the top (Figure 1H).

### Luciferase assay

For BMP reporter assay (Figure 2B), Renilla DNA (pRL; Promega, 25 pg) and BMP reporter DNA (50 pg) were co-injected into both blastomeres at two-cell stage with or without 15 ng *fhl3* MO. Twenty DLMZ explants were dissected at stage 10.5, grown in ¾ NAM, and lysed in 60 µl of Passive lysis buffer (Promega) at stage 12.5.

For WNT reporter assay in ectoderm/mesoderm recombinants (Figure 2F), Renilla DNA (25 pg) and WNT reporter DNA (TOPFLASH, 50 pg) were co-injected into both blastomeres at two-cell stage. Animal cap (ectoderm) explants were dissected at stage 9 and juxtaposed to control or Fhl3 depleted DLMZ. 15 recombinant explants were lysed in 60 µl Passive lysis buffer at stage 18.

For BMP and WNT reporter assays in iNC (Figure 3C), Renilla DNA (25 pg), *pax3-GR* mRNA (35 pg) and *zic1-GR* mRNA (72 pg) were co-injected with either BMP reporter DNA (50 pg) or WNT reporter DNA (50 pg) into both blastomeres at two-cell stage with or without 15 ng of *fhl3* MO. Twenty animal cap (ectoderm) explants were dissected out from stage 9 embryos and lysed in 60 µl of Passive lysis buffer at stage 18.

In all assays, Firefly and Renilla luciferase activities were measured using the Dual-Glo Luciferase Assay System (Promega).

### qRT-PCR

Total RNA was extracted using Phenol/Chloroform/Isoamyl Alcohol from *Xenopus laevis* embryos or tissue explants at late neurula stage (stage 18). We used M-MLV reverse transcriptase for reverse transcription and SYBR Green mix (BIO-RAD) for quantitative RT-PCR. Results were normalized against the reference genes *odc* and *ef1a*. For primers sequences see Table S1.

### Chromatin immunoprecipitation (ChIP)

Embryos were injected in both blastomeres at two-cell stage with mRNA encoding FLAG-Smad1 (200 pg) and FLAG-Fhl3 cDNA separately (200 pg) or FLAG-Fhl3 cDNA with noggin mRNA (25 pg). Injected embryos were collected at late gastrula stage (stage 12, 100 embryos/sample) and processed according to (Wills et al., 2014). Sonication was performed using a 3.2 mm tapered microtip for 13 mm horn (Branson 450 sonicator). Anti-FLAG antibodies (42 μg) coupled to magnetic beads were added to the chromatin. Quantitative PCR was performed with immunoprecipitated fragmented chromatin using *wnt8* and *ventx1.1* promoter region primers. qPCR results were normalized to *ef1a* and input. Primers and FLAG antibody are listed in Table S1.

### Western blot detection of phospho Smads

10 to 15 DLMZ explants were dissected at stage 10.5, lysed when sibling embryos reached stage 12.5 in presence of phosphatase inhibitor (PhosphoSTOP, Roche) and protease inhibitors (Sigma) and analyzed by standard western blotting. ECL signal was quantified using ImageJ. Antibodies are listed in table S1.

### Statistical analysis, imaging and image processing

All experiments were performed at least in biological triplicates (except the rescue experiment shown in Figure 2D, the Fhl3-FLAG + noggin ChIP experiment (Figure 4C) and *axin2* and *wnt8* rescue experiment in Figure 4D, which were performed in biological duplicates). Each dot on the graphs represents one independent experiment. Examples of embryos with the most frequent phenotypes are shown. We used Prism for statistical analysis, error bars represent S.E.M. and Student’s *t*-test was used to determine statistical significance (*p<0.05; **p<0.01; ***p<0.001, ns: not significant). Images were taken using a Lumar V12 Binocular microscope equipped with bright field or fluorescent filters and color camera (Zeiss), and were processed using Photoshop.

---

**KEY RESOURCES TABLE.**
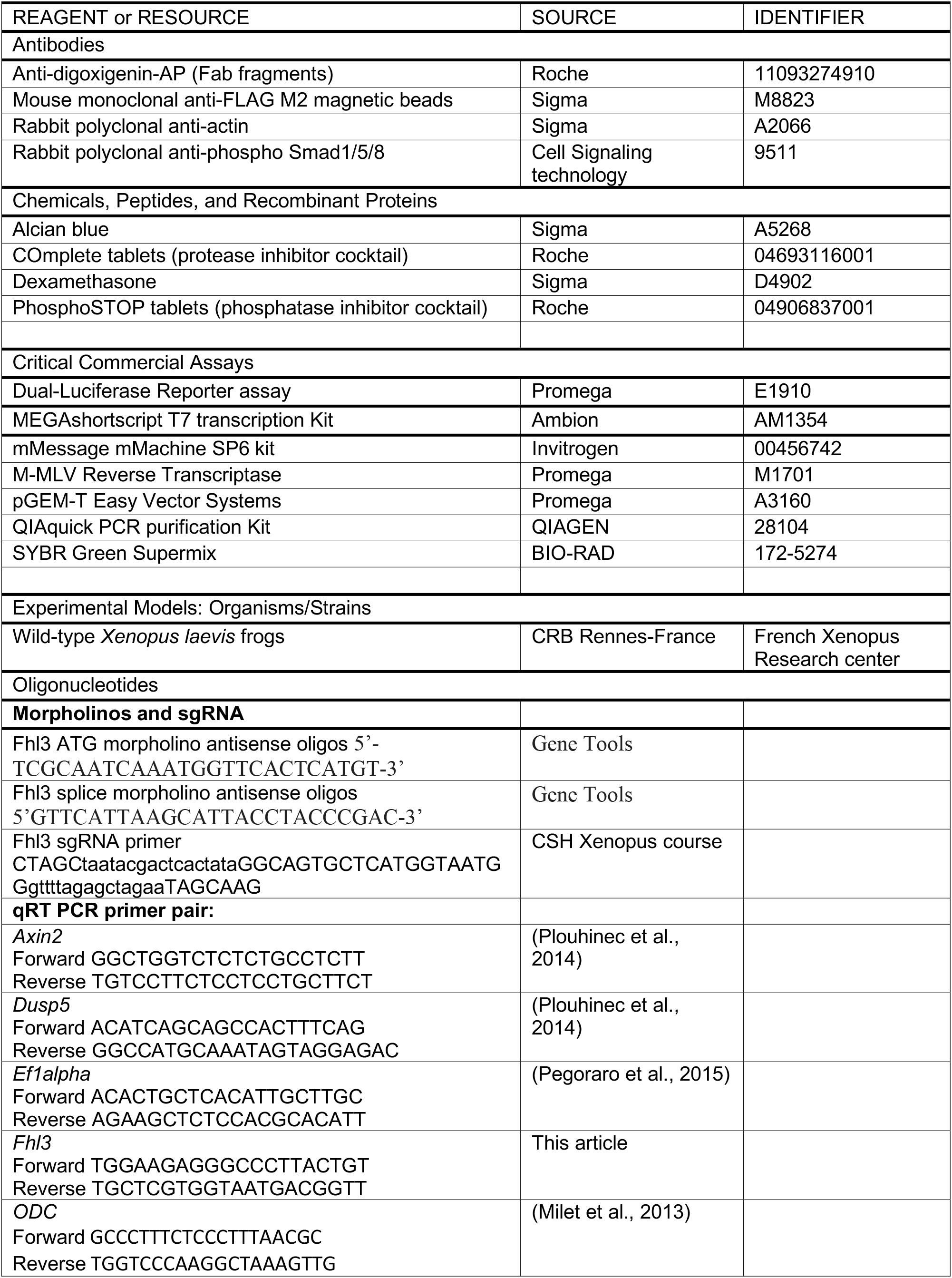
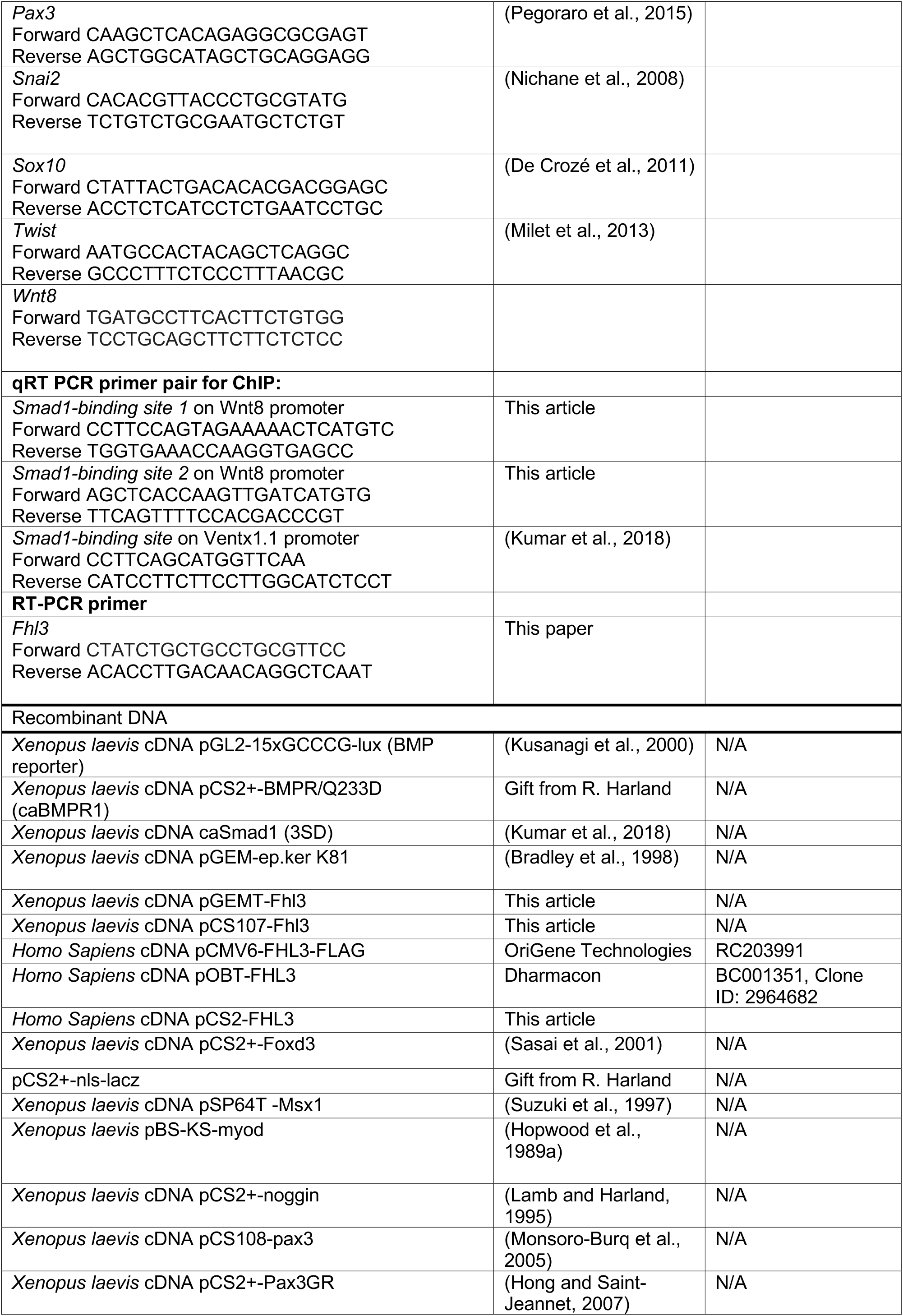
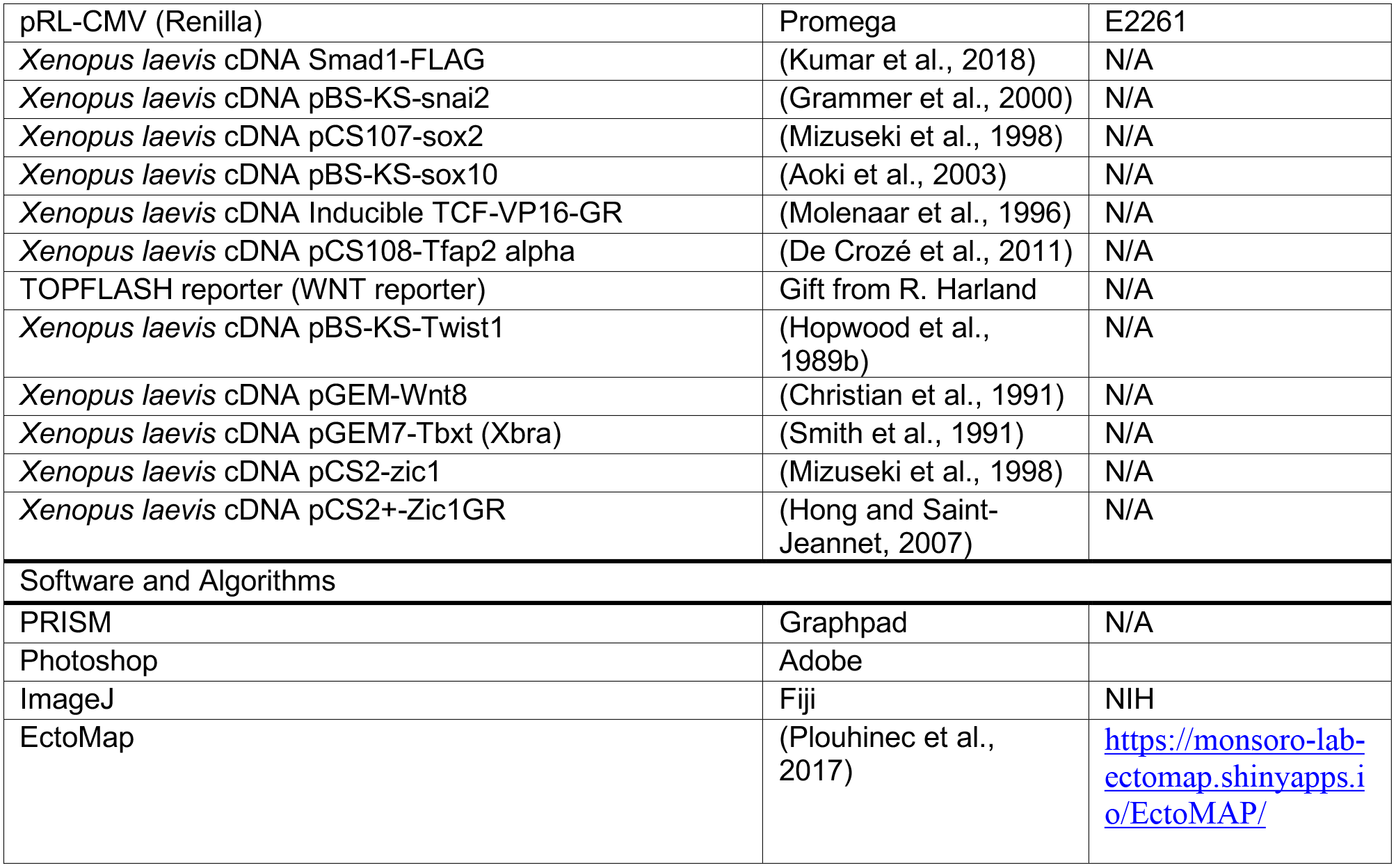

